# Compositionally aware estimation of cross-correlations for microbiome data

**DOI:** 10.1101/2023.05.22.538214

**Authors:** Ib Thorsgaard Jensen, Luc Janss, Simona Radutoiu, Rasmus Waagepetersen

## Abstract

In the field of microbiome studies, it is of interest to infer correlations between abundances of different microbes (here referred to as operational taxonomic units, OTUs). Several methods taking the compositional nature of the sequencing data into account exist. However, these methods cannot infer correlations between OTU abundances and other variables. In this paper we introduce the methods SparCEV (Sparse Correlations with External Variables) and SparXCC (Sparse Cross-Correlations between Compositional data) for quantifying correlations between OTU abundances and either continuous phenotypic variables or components of other compositional datasets, such as transcriptomic data. We compare these new methods to empirical Pearson cross-correlations after applying naive transformations of the data (log and log-TSS). Additionally, we test the centered log ratio transformation (CLR) and the variance stabilising transformation (VST). We find that CLR and VST outperform naive transformations, except when the correlation matrix is dense. For large numbers of OTUs, SparCEV and SparXCC perform similarly to CLR and VST. SparCEV outperforms all other tested methods when the number of OTUs is small (less than 100). SparXCC outperforms all tested methods when at least one of the compositional datasets has few variables (less than 50), and more so when both datasets have few variables.

**Author summary:** Sequencing data of the microbiome posses a unique and challenging structure that renders many standard statistical tools invalid. Features such as compositionality and sparsity complicates statistical analysis, and as a result, specialized tools are needed. Practitioners have long been interested in the construction of correlation networks within the microbiome, and several methods for accomplishing this exist. However, less attention has been paid to the estimation of cross-correlations between microbial abundances and other variables (such as gene expression data or environmental and phenotypic variables). Here, we introduce novel approaches, SparCEV and SparXCC, for inferring such cross-correlations, and compare these to transformation-based approaches, namely log, log-TSS, CLR and VST. In some cases, SparCEV and SparXCC yield superior results, while in other cases, a simpler transformation-based approach suffices. The methods are used to study cross-correlations between bacterial abundances in the skin microbiome and the severity of atopic dermatitis, as well as cross-correlations between fungal and bacterial OTUs in the root microbiome of the legume *Lotus japonicus*.

## Introduction

Sequencing data are ubiquitous in modern biology [1]. For example, RNA-seq data have been used to identify genes associated with clinical outcomes of cancer patients [2], for human disease profiling [3], and to identify genes with possible links to Rett Syndrome [4]. Microbiome data have drawn much attention in recent years, particularly regarding the human gut microbiome. Composition of the human gut microbiome has been shown to be associated with several aspects of human health, such as obesity [5] and metabolic disorders [6]. More recently, the integration of microbiome data with other omics data has received increasing interest [7–11].

Data from sequencing technologies pose many specific challenges. They produce count data with technical noise, which makes results from rare features difficult to interpret. Additionally, they are compositional, meaning that the observed variables are components of an arbitrary total.

Within the field of microbiome studies, several methods have been proposed to infer interactions between microbial abundances. These include Local Similarity Analysis (LSA) [12], which finds non-linear relationships through time using time series data; Sparse Compositional Correlations (SparCC), which infers correlations based on compositional data [13]; and Sparse Inverse Covariance Estimation for Ecological Association Inference (SPIEC-EASI), which infers relations through graphical models [14]. In this paper, we focus on the estimation of correlations.

When considering correlations in the context of compositional data, there are essentially three cases of interest: A) correlations between features of the same compositional dataset, B) cross-correlations between features of a compositional dataset and non-compositional variables, and C) cross-correlations between two compositional datasets. Correlations between bacterial abundances in a microbiome is an example of case A. An example of case B is cross-correlations between gut microbes and clinical features of patients [15], and an example of case C is cross-correlations between microbial abundances and gene expression levels from RNA-seq data [7]. For an overview of these cases, see Table 1.

**Table 1.** Cases A, B, and C along with explanations and and overview of applicable methods.

| Case | Correlations | Methods |
| --- | --- | --- |
| A | Within a composition | SparCC<br>SPIEC-EASI<br>LSA<br>Pearson |
| B | Between a composition and an external variable | SparCEV<br>Pearson |
| C | Between two compositions | SparXCC<br>SPIEC-EASI<br>mmVec<br>Pearson |

The methods mentioned above all operate in case A. More recently, case C has gained more interest, with new methods being developed. For example, SPIEC-EASI has been extended to infer interactions between variables from two compositional datasets [16]. Like the original SPIEC-EASI, pairs of variables that are conditionally independent are identified by estimating the precision matrix using a penalized estimation scheme to enforce sparsity. The method mmVec [17] is designed for identifying interactions between OTU abundances and metabolite concentrations. This method employs a neural network to estimate the probability of observing a metabolite, given that a specific OTU is observed. It was shown to perform with similar accuracy as the extended SPIEC-EASI and to outperform correlation-based procedures. However, the correlations were estimated using a flawed methodology, where the centered log-ratio (CLR) transformation was applied to both datasets simultaneously, rather than separately, thus biasing the results. This was noted in a response from Quinn and Erb [18], who showed that when the CLR-transformation was applied appropriately, correlation-based methods outperform both mmVec and SPIEC-EASI in the setup examined by Morton et al. [17]. However, in a counter-response Morton et al. showed that it was possible to construct scenarios, where mmVec outcompeted all alternatives. Futhermore they show that mmVec is better suited to handle sparse data than correlation-based methods [19].

In this paper, we focus on inferring cross-correlations in cases B and C. Inspired by SparCC, we introduce two novel compositionally aware methods, SparCEV (Sparse Correlations with External Variables) and SparXCC (Sparse Cross-Correlations between Compositional data). Using simulation studies, we compare these methods to Pearson cross-correlations applied to various transformations of the data. Theoretical comparisons of transformation-based methods and derivations of new methods are given in the supplementary material.

## Materials and methods

### Modelling Sequencing Data

Let *a*_*i*_ denote the absolute abundance of OTU 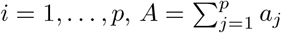and *r*_*i*_ = *a*_*i*_*/A* the relative abundance. The aim of this paper is to estimate the correlation between log *a*_*i*_ and other log transformed variables. However, we only have access to observed read counts, denoted *x*_*i*_ for OTU *i*. To theoretically compare the different strategies and to develop new methods, we adopt a simplified modelling framework, where

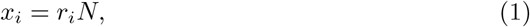

where 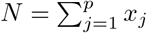 denotes the library size. We compare the methods considered using simulations from models that are more complex and realistic than (1), see Section Simulation models. In these models, *x*_*i*_ given (*r*_*i*_, *N*) is not fixed. We use the term *technical variance* for the variance of *x*_*i*_ given (*r*_*i*_, *N*) and the term *biological variance* for the variance of *r*_*i*_. The more realistic models are, however, intractable for theoretical analysis.

All tested methods but one require log-transformation of the *x*_*i*_s, which is problematic if *x*_*i*_ = 0 is observed. As a remedy, we add 1 to all read counts prior to log-transformation of the data (the pseudo-count method).

### Existing Strategies for Cross-correlation Estimation

#### Naive transformation

We use the term *naive transformation* to refer to any transformation that does not take the compositional nature of the data into account. Naive transformations considered in this paper are log and log total sum scaling (TSS). Theoretically, naive transformations do not adequately account for the compositional structure of the data (see S1 Text). Nonetheless, it remains a common practice to apply these transformations [20] or no transformations at all [15, 21–25]. As a result, any method that outperform naive transformations would constitute an improvement relative to common practice.

#### Adapted Transformations

We use the term *adapted transformations* to refer to transformations that are adapted to the particular structure of the data beyond differing library sizes. In this paper, we consider the centered log-ratio (CLR) and the variance-stabilising transformation (VST). See Table 2 for definitions of all transformations (naive and adapted) considered in this paper. Some common transformations, such as trimmed M-means (TMM) [26], DESeq’s median-based transformation [27], and upper-quartile transformation (UQ) [28], are not included, since they do not correct for within-replicate biases and are thus not applicable for correlation estimation.

**Table 2.** An overview of the transformations used to assess cross-correlations. In VST, *k* is a replicate index.

| Transformation | Expression | Interpretation |
| --- | --- | --- |
| log | $\log x_i$ | Log-transformed observed read counts |
| log-TSS | $\log \text{TSS}(x_i) = \log \left( \frac{x_i}{\sum_{j=1}^p x_j} \right)$ | The log of estimated relative abundances |
| CLR | $\text{CLR}(x_i) = \log x_i - \frac{1}{p} \sum_{j=1}^p \log x_j$ | Abundances relative to the average abundance |
| VST | $\text{VST}(x_{ik}) = \int_0^{x_{ik}/\hat{s}_k} f(\mu) d\mu$,<br>where $f(\mu) = \mu^2 \hat{a}_0 + \mu(1 + \hat{a}_1)$ | Removal of mean-variance relationship |

VST makes use of the DESeq modelling framework [27]. Specifically, it assumes that *x*_*ik*_ *∼* NB(*µ*_*ik*_, *φ*_*i*_), where *µ*_*ik*_ = *s*_*k*_*λ*_*i*_ and *φ*_*i*_ = *a*_0_ + *a*_1_*/λ*_*i*_. The estimates 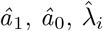and *ŝ*_*k*_ are obtained using the DESeq2 estimation procedures [27, 29]. Here, *k* denotes the index of the biological replicate. VST requires at least one feature without any zero counts. This may be violated for data with small *p* which is frequently the case for our simulated microbiome data. We therefore only consider the VST transformation in case C for simulated gene expression data while applying CLR to the microbiome data.

For convenience, we use the term CLR for the method where empirical Pearson cross-correlations are applied to CLR-transformed data, and likewise for log, log-TSS, and VST.

#### Theoretical assessment of transformations

We examine theoretically whether empirical Pearson cross-correlations combined with the transformations presented in Table 2 are likely to yield good approximations of cross-correlations. We focus on case B, since it is simpler and the results in case C are analogous. We seek an approximation of ℂorr [log *a*_*i*_, *b*], where *b* is a non-compositional variable, here referred to as a *phenotypic variable*. By (1) and Table 2 we have,

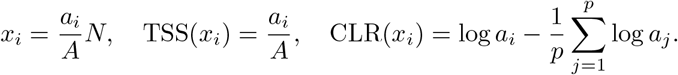

By definition,

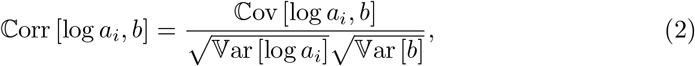

and we require good approximations of the numerator and the denominator. Since *b* is not compositional, 𝕍ar [*b*] can easily be estimated without the need for approximation. According to our derivations in S1 Text, log, log-TSS, and CLR all lead to reasonable approximations of the covariance ℂov [log *a*_*i*_, *b*] under the model in (1) given appropriate assumptions. Furthermore, we show that

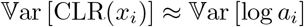

when *p* is large. However, analogous results do not hold for log and log-TSS, demonstrating that naive transformations are not sufficient. Summing up, CLR can yield good approximations under the model in (1) with the following assumptions:

(Bi) 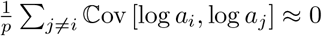for all *i*

(Bii) ^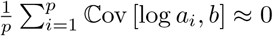^ for all *i*

(Biii) The number of OTUs, *p*, is large.

In case C, we we need (Bi), (Biii), and the additional assumptions

(Ci) 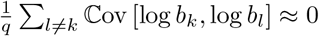for all *k*_*q*_

(Cii) 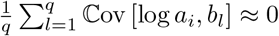 for all *i*_*p*_

(Ciii) 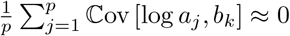 for all *k*

(Civ) The number of genes, *q*, is large

Conditions (Bi), (Bii) and (Ci)-(Ciii) hold if the correlation matrix is sparse; thus, following the language of Friedman and Alm [13], we refer to these as *sparsity assumptions*. This is a slight abuse of terminology, since these conditions may also hold if all rows of the correlation matrix contain entries whose distributions are symmetric around zero, even though such a matrix is not sparse.

#### Compositionally Aware Methods

Inspired by SparCC, we introduce compositionally aware methods for cases B and C. In case B, we assume the same sparsity condition as for CLR in the previous section. We then show in S1 Text that

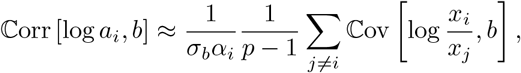

where 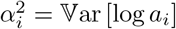 can be estimated by SparCC and 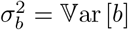 can be estimated in a standard fashion. In contrast to CLR this method only requires conditions (Bi) and (Bii), but not (Biii). Therefore, it is likely preferable when *p* is small. We name this method ***Spar****se* ***C****orrelations of* ***E****xternal* ***V****ariables* (SparCEV).

In Case C, we let *b*_*k*_, *k* = 1, …, *q*, denote the gene expression level of the *k* th gene, 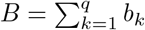 and *M* the library size. Similar to (1), we assume the model

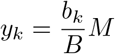

for the observed gene expression level. In S1 Text, we obtain the relation

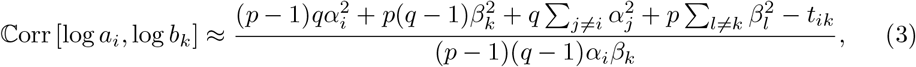

where

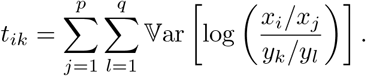

The parameters 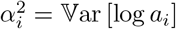 and 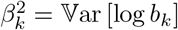 can be approximated by applying SparCC to the microbiome and gene expression datasets individually, and the variances in *t*_*ik*_ can be estimated in a standard fashion. Details regarding efficient computation of the *t*_*ik*_s are given in the S1 Text. As with CLR, we need the assumptions (Bi) and (Ci)-(Ciii), but unlike CLR, we do not need (Biii) and (Civ). We refer to this method as ***Spar****se Cross-****C****orrelations of* ***C****ompositional data* (SparXCC, where “X” represents “cross”).

#### Simulation models

We adopt the parametric model employed by *SparseDOSSA2* [30], adapting the methodology slightly to handle cases B and C. A simulated dataset contains *n≥* 1 replicates, where *n* is the number of microbiome samples sequenced. The individual simulated variables (e.g. abundances or gene expression levels) are characterized by the mean, *µ*_*i*_, variance, 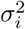, and zero-probability, *π*_*i*_. The parameter *π*_*i*_ reflects the probability that OTU *i* is absent from a given replicate. We refer to this as a *biological zero*. The correlation between variables is characterized by the correlation matrix *R*. We simulate *p* OTU abundances, *a*_1_, …, *a*_*p*_ and *q* other variables, *b*_1_, …, *b*_*q*_. In case B, the latter *q* variables are non-compositional, typically *q* = 1, and we take *π*_*p*+*k*_ = 0,*k* = 1, …, *q*, so that biological zeros do not occur for the *b*_*k*_s. In case C, *q >* 1 and the *b*_*k*_s are compositional. The library size *N*^*a*^ is simulated from a log normal distribution with parameters *µ*_*a*_ and 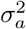.

The simulation algorithm is given in the following steps. For ease of presentation, we present the case where *n* = 1, but when *n≥* 1 the steps would simply be repeated *n* times.

1. Simulate the *p* + *q*-dimensional variable *g ∼* N(0, *R*).
2. Define the variables *Z*_*i*_ for *i* = 1, …, *p* + *q* such that *Z*_*i*_ = 0 if *g*_*i*_ *<* Φ^*−*1^(*π*_*i*_) and 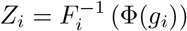 otherwise, where Φ is the standard normal cumulative distribution function (cdf) and 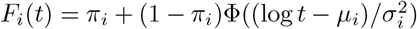 is the cdf of a zero-inflated log-Gaussian distribution with parameters 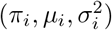. We now have

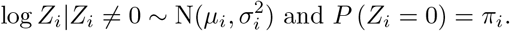
3. Set *a*_*i*_ = *Z*_*i*_ for *i* = 1, …, *p* as the absolute OTU abundances. In case C, *b*_*j*_ = *Z*_*j*+*p*_ for the absolute gene expression levels. In case B, we let *b*_*j*_ = log *Z*_*j*+*p*_ for the non-compositional phenotypic variables.
4. Set 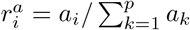 as the relative abundances of the OTUs, and in case C, set 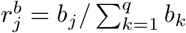 as the relative expression levels.
5. Simulate 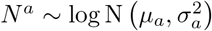 and let ⌈*N*^*a*^⌉ be the library size.
6. Simulate the vector, *x* = (*x*_1_, …, *x*_*p*_)^*T*^, of observed read counts of OTU 1,…, *p* as *x ∼* Multinom 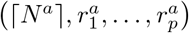
7. In case C, simulate 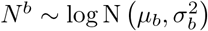 and let ⌈*N*^*b*^⌉ be the library size.
8. In case C, simulate the vector, *y* = (*y*_1_, …, *y*_*q*_)^*T*^, of observed read counts of gene *j* = 1, …, *q* as *y ∼* Multinom 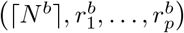.

In case B, steps 7–8 are skipped. Excluding the simulation of the *b*_*j*_s, steps 1–6 in the above procedure are identical to the procedure employed by SparseDOSSA2. The above simulation scheme differs from the model (1) by the multinomial noise generated in steps 6 and 8 where the multinomial model is a simplistic representation of the randomness generated in the sequencing procedure. The correlation ℂ orr [log *a*_*i*_, log *a*_*j*_] agrees with *R*_*ij*_ when *π*_*i*_ = *π*_*j*_ = 0. This does not hold in the presence of biological zeros, *π*_*i*_ *>* 0 or *π*_*j*_ *>* 0, in which case log *a*_*i*_ or log *a*_*j*_ may not even be well defined. We nevertheless use *R* as a ground truth for comparison with our estimates, including the case of biological zeros. In that way, the presence of biological zeros is considered a source of noise relative to our method in addition to the multinomial noise.

The correlation matrix *R* is constructed using two methods described in S1 Text. The first is called the cluster method, and it works by assigning a portion of the OTUs to a “cluster”. All OTUs in the cluster are correlated to each other with the same correlation coefficient and uncorrelated to every other OTU. All OTUs outside the cluster are also uncorrelated with each other. In case C, a similar portion of the genes are also assigned to the cluster, and in case B, all non-compositional variables are also assigned to the cluster. This gives us a high degree of control over the degree of sparsity and the strength of the correlations. The second method is called the loadings method, and it results in a correlation matrix without exact zero entries but where most variables are only weakly correlated, with a relatively small proportion of highly correlated variable pairs. The loadings method most likely results in more realistic systems than the cluster method.

Throughout the simulations in this paper, we simulate *n* = 50 replicates. In many practical settings, *n* is considerably lower than that. However, for the purposes of the present simulation study, it is important that we can detect biases in the estimators. If the bias is small relative to the variance of the estimator, it may be difficult to detect in a simulation study. Since the variance of an estimator increases as *n* decreases, it is counter-productive to perform simulation studies with small *n*. In other words, we construct a situation where the main bottleneck to producing accurate results is the chosen method, not the size of the dataset.

#### Selecting Parameter Values

In the simulations results shown in Figs 1 and 4, the log-scale parameters, *µ*_*i*_ and 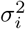 and zero-probabilities, *π*_*i*_, are chosen using a real dataset as a template. We estimate the mean, *µ*_*ri*_, and variance, 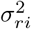 of the observed read counts for *i* = 1, …, *p*. We then choose *µ*_*i*_ and 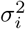 such that the simulated variables have mean *µ*_*ri*_ and variance 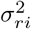, on the linear scale. By the properties of the log-normal distribution, the means and variances are related by

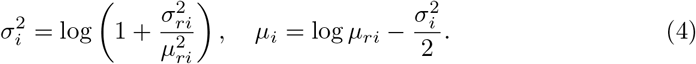

**Fig 1.**
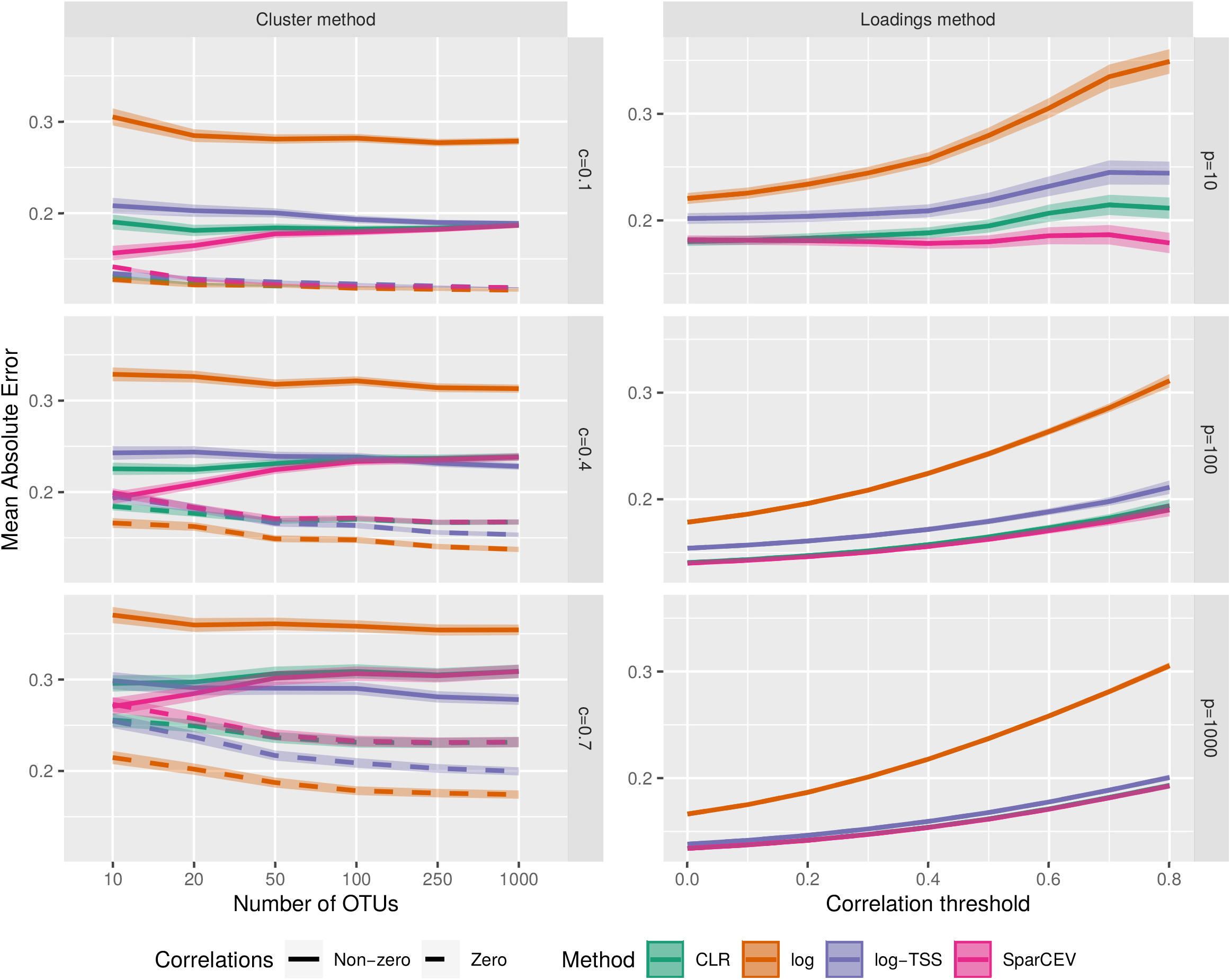
MAE of different cross-correlation methods for correlation matrices generated by the cluster method (left column) and the loadings method (right column). For the cluster method, different *p* (number of OTUs) and *c* (the proportion of OTUs in a cluster) are used. For the loadings method, threshold values *t* = 0, 0.1, …, 0.8 and different *p* are used. The lines show the mean accuracy, and the edges of the envelopes show ±1.96 standard errors (SE). The results are based on 1000simulated datasets where each simulated dataset has 50 replicates.

**Fig 2.**
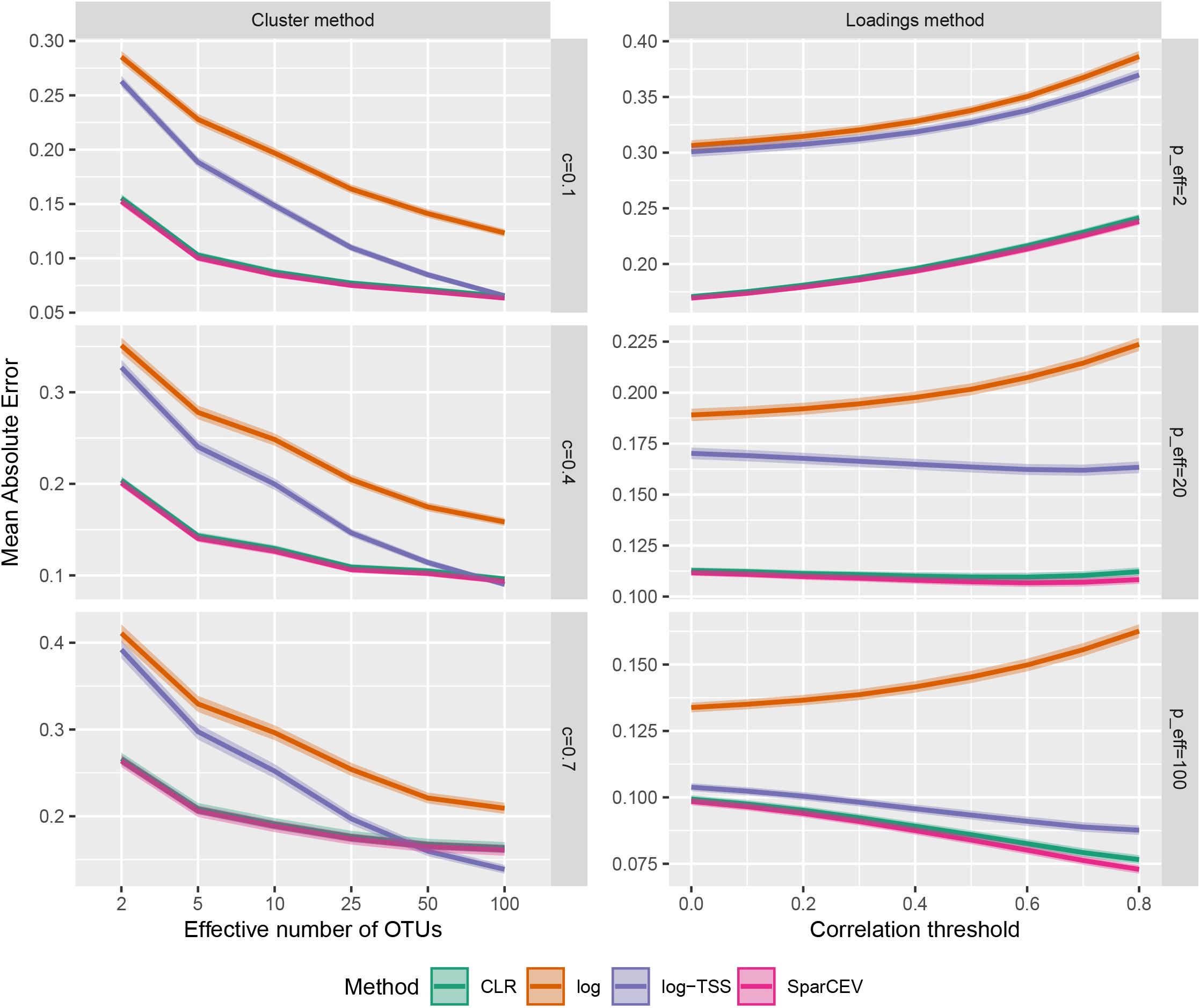
MAE of different cross-correlation methods for correlation matrices generated by the cluster method (left column) and the loadings method (right column). For the cluster method, different *p*_eff_ (effective number of OTUs) and *c* (the proportion of OTUs in a cluster) are used. For the loadings method, threshold values *t* = 0, 0.1, …, 0.8 and different *p*_eff_ are used. The lines show the mean accuracy, and the edges of the envelopes show ±1.96 SE. The results are based on 1000simulated datasets where each simulated dataset has 50 replicates.

**Fig 3.**
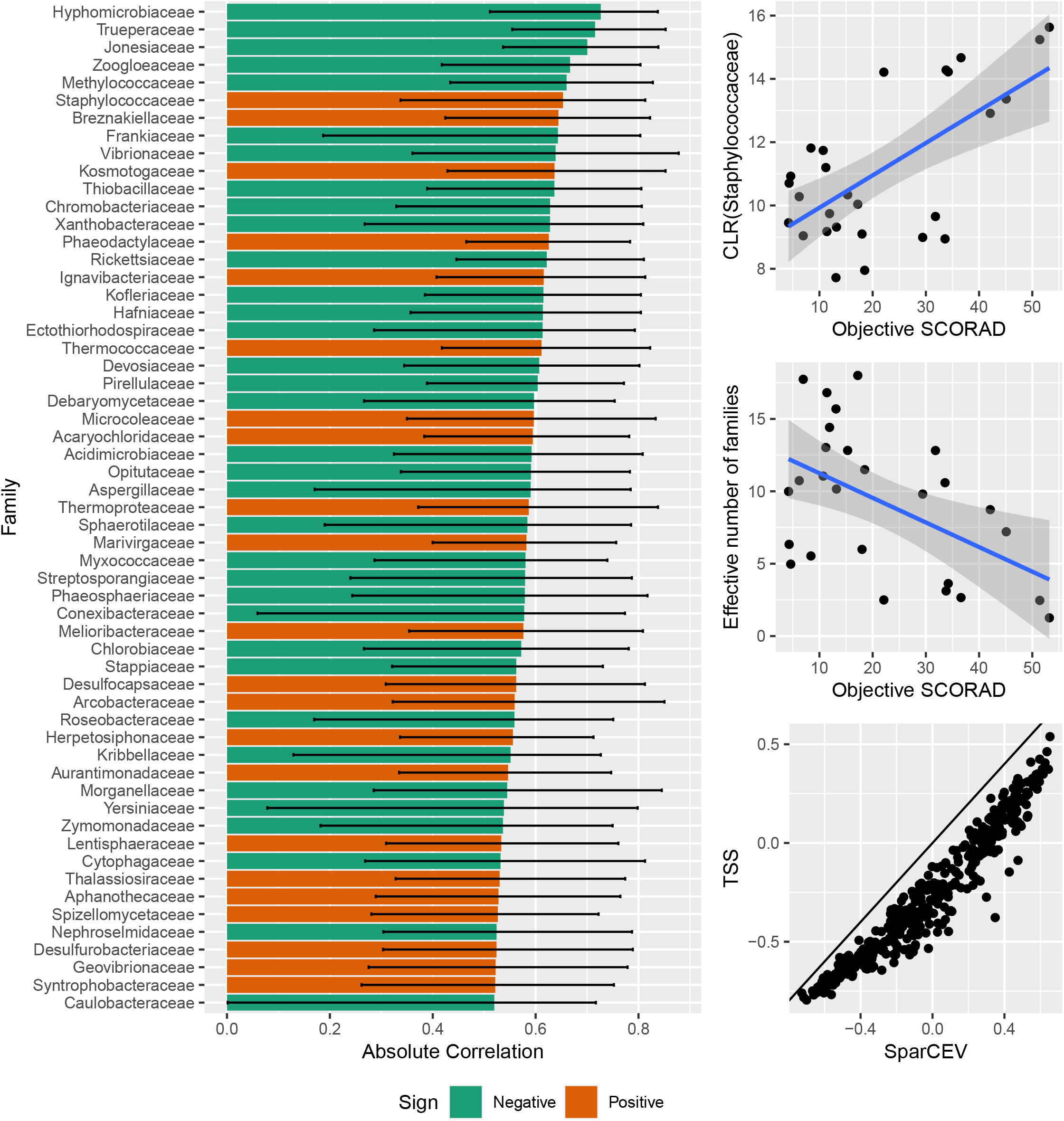
Results from a correlation analysis on atopic dermatitis data from Byrd et al. [31]. **A**: All correlations exceeding the permutation threshold of 0.52 with color according to the sign of the correlation and with error bars given by the empirical bootstrap 95%-confidence interval. **B**: Scatter plot between the CLR-transformed abundance of *Staphylococcaceae* and the objective SCORAD. The blue line is derived from a smooth line fitted to the data with error bars derived from the standard deviation. **C**: Scatter plot between the effective number of families and the objective SCORAD. **D**: Scatter plot between the estimated correlation using log-TSS and SparCEV. The straight line has slope 1 and intercept 0.

**Fig 4.**
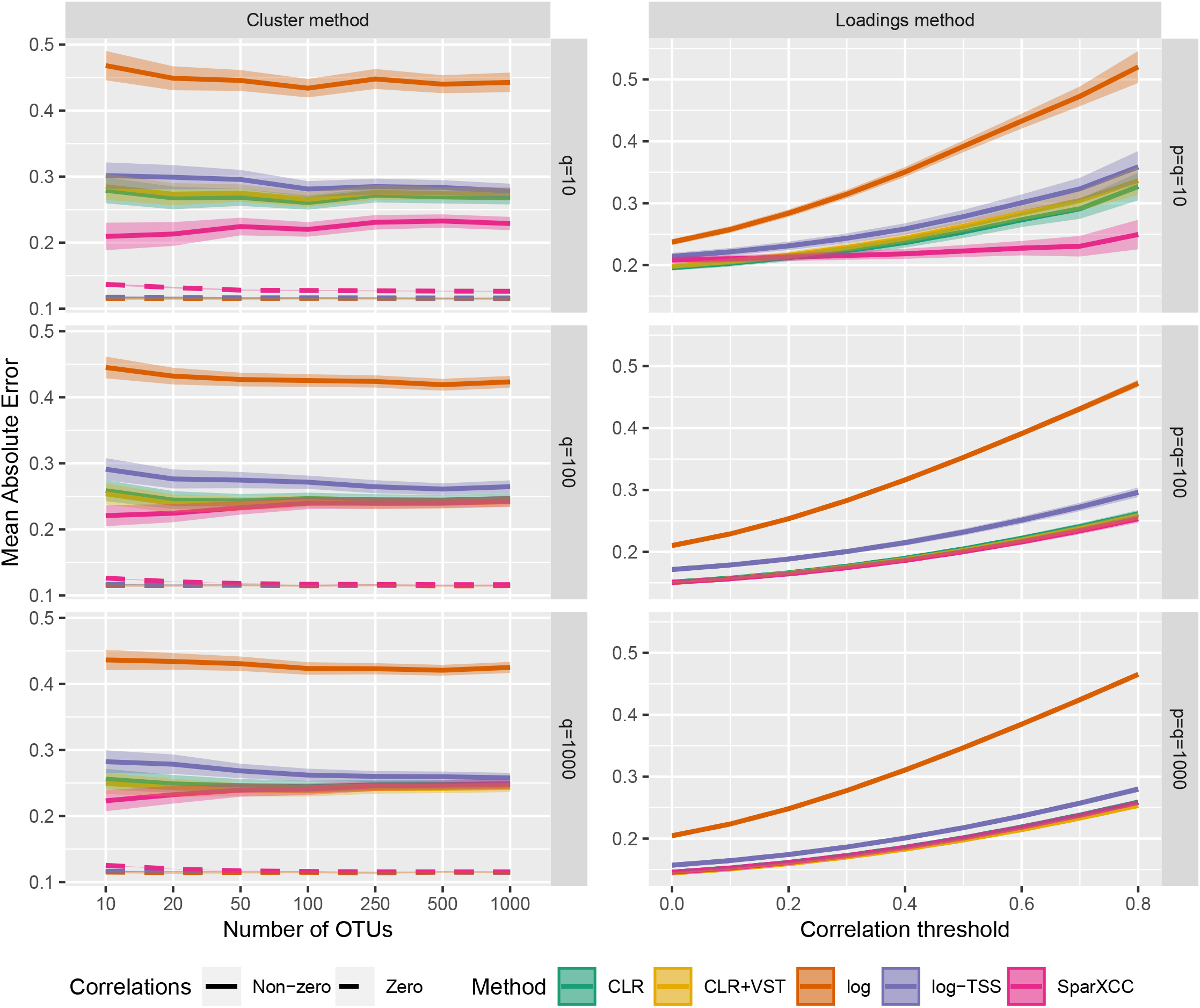
MAE of different cross-correlation methods for correlation matrices generated by the cluster method (left column) and the loadings method (right column) in case C. For the cluster method, different *p* (number of OTUs) and *q* (number of genes) are used. For the loadings method, threshold values *t* = 0, 0.1, …, 0.8 and different *p* and *q* are used. The lines show the mean accuracy, and the edges of the envelopes show ±1.96 standard errors (SE). The results are based on 500 simulateddatasets where each simulated dataset has 50 replicates.

The parameters *π*_*i*_ are set to half the proportion of zeros for the *i*th variable, with the assumption that half of the zeros are biological and the other half are technical. In case B, the parameters of the phenotype variable are somewhat arbitrarily chosen so that it has a mean of 30 and a variance of 1.

In Fig 2, considered later, we examine the impact of diversity on the accuracy of the correlation estimation methods. We measure diversity using the *effective number of OTUs, p*_eff_. We have *p*_eff_ = e^*H*^, where 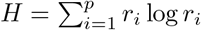 is the entropy or *Shannon index*. The quantity *p*_eff_ can be interpreted as the minimal number of OTUs such that a replicate has entropy *H*. This occurs when all OTUs are equally abundant. We choose 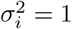and *π*_*i*_ = 0 for *i* = 1, …, *p* and *µ*_*i*_ in such a way that we get a specific value of *p*_eff_ in expectation. This is accomplished by selecting the linear-scale mean relative abundances 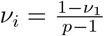 for *i ≥* 2 and obtaining *ν*_1_ by solving the equation

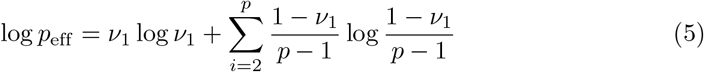

for *ν*_1_ given a choice of *p*_eff_. We then choose an arbitrary value for the microbial load, say 1000, and set *µ*_*ri*_ = 1000*ν*_*i*_. Finally, the *µ*_*i*_ are obtained from the right part of (4).

#### Methods Assessment

We might assess the accuracy of correlation estimates 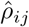 by comparing the true correlations to the estimated correlations by computing, for example, the mean absolute error (MAE). However, suppose we use the estimate 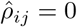 for all *i, j* and pick, for example, *c* = 0.05 and *ρ* = 0.75 for the cluster method. Then, even though non-zero correlations are not well estimated, the MAE, *cρ* = 0.0375, is quite low. Thus, we separately consider the MAE of the pairs whose true correlation is zero and the MAE of the pairs whose true correlation is non-zero. In case of the loadings method, no correlations are exactly zero, so we then instead assess the MAE of pairs whose true correlation-coefficient exceeds the thresholds 0, 0.1, …, 0.8.

In summary, for the cluster method, we use the criteria

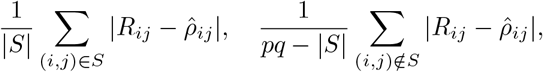

where *S* ={ (*i, j*) ∈*A*_*pq*_ *R*_*ij*_ ≠ 0 }and *A*_*pq*_ ={ (*i, j*) ∈ℕ^2^ |0 ≤*i≤ p, p < j ≤p* + *q }*. For the loadings method, we use the criteria

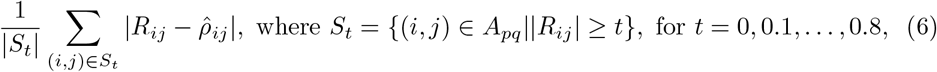

where *t* = 0 corresponds to the overall MAE.

#### Discriminating between Correlated and Uncorrelated pairs

Since sparsity is only an approximate assumption, any test-statistic used to derive *p*-values is likely to be biased. This is exacerbated by the technical noise, which has particularly high impact for low-abundance OTUs. We shall not attempt to remedy these challenges here. Instead, we choose a dynamic threshold based on the data. Pairs whose estimated absolute correlation exceeds this threshold are considered the most likely candidates for genuinely correlated pairs. The threshold is derived in the following way. Let *X* and *Y* be the two datasets under study (in case C, *Y* is compositional). Permute each dataset separately. This breaks all cross-correlation, but not the correlations within each dataset. Let *S* _perm_ be the set of cross-correlation estimates obtained from the permuted data and let *m* be the 100(1-1/(*pq*))-percentile of *S* _perm_.

We might use *m* as a threshold, since almost all correlations should be smaller than *m*. However, since all correlations are broken under the permutation, *m* may be too low for real data where not all correlations are zero. More specifically,

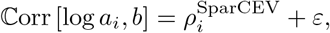

where 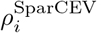 is the SparCEV estimate and where

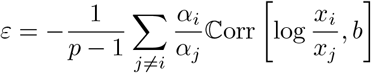

in practice may differ from zero due to violation of sparsity condition (Bii). In practice, *ε* may be small relative to the noise of 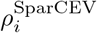 if *n* is sufficiently low. However, as *n* grows, the noise of 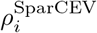 shrinks, and the relative impact of *ε* grows. This may lead to an unacceptable amount of false discoveries, as we see in S10 Fig. To remedy this, we add an additional user-specified parameter, *t*, where only cross-correlations above *t* are considered to be of interest. We then use *m*^*∗*^ = max *{m, t}* as the final threshold. By S9 Fig and S10 Fig, this sufficiently controls the false discovery rate when *n* is large.

#### Data Availability

All data used in this paper can be found at https://github.com/IbTJensen/Microbiome-Cross-correlations/. The raw sequencing data from Byrd et al. [31] can be found in NCBI Bioproject 46333, and the OTU table was originally obtained from Morton et al. [32] at https://github.com/knightlab-analyses/reference-frames. The raw sequencing data from Thiergart et al. [20] can be found at the European Nucleotide Archive (ENA). The 16S dataset has project accession no. PRJEB34100, and the ITS dataset has project accession no. PRJEB34099. The OTU tables was originally obtained at https://github.com/ththi/Lotus-Symbiosis.

#### Implementation and Code Availability

Illustrations are produced using ggplot2 version 3.4.1 [33], ggpubr version 0.6.0 [34], and GGally version 2.1.2 [35]. The VST transformation was performed using DESeq2 version 1.34.0 [29], hypothesis testing on cross-correlations were carried out using psych version 2.2.9 [36], and the SPIEC-EASI networks were estimated using the package SpiecEasi version 1.1.2 [37]. The scripts used for the simulations and the code for SparCEV and SparXCC can be found at https://github.com/IbTJensen/Microbiome-Cross-correlations/.

## Results

In this section, we compare the different estimation methods on simulated datasets with the correlation matrices constructed using the cluster and the loadings methods.

### Case B

Fig 1 shows the performance of the different correlation estimation methods, with correlation matrices generated by both the cluster and the loadings method. All MAEs are computed as means over 1000 simulated datasets, with *n* = 50 replicates. For the cluster method *ρ* = 0.75 and for the loadings method *k* = 5. With both correlation generation methods, poor results are obtained when only the log-transformation is applied, and all other methods yield better results. For the cluster method, CLR and SparCEV outperform log-TSS when *c* = 0.1, and SparCEV outperforms CLR when *p* is small. When *c* = 0.4, log-TSS performs the same or better than CLR and SparCEV for*p ≥*100. For *c* = 0.7, 70% of pairs are correlated, and thus the sparsity assumption is severely violated. As expected, this is a substantial obstacle to accurate estimation, especially for CLR and SparCEV. In fact, log-TSS performs similarly or better than both, except when *p* = 10, where SparCEV still has a slight edge. For the loadings method, SparCEV outperforms all alternatives when *p* = 10. For *p* = 100, the difference between SparCEV and CLR is negligible, but both outperform log-TSS. When *p* = 1000, SparCEV and CLR perform practically identically, and they only outperform log-TSS at higher thresholds and only by a small margin. For both the cluster and the loadings methods, the difference between CLR and SparCEV shrinks as *p* increases, which is in agreement with the theory presented in S1 Text.

Caption for Fig 1: MAE of different cross-correlation methods for correlation matrices generated by the cluster method (left column) and the loadings method (right column). For the cluster method, different *p* (number of OTUs) and *c* (the proportion of OTUs in a cluster) are used. For the loadings method, threshold values *t* = 0, 0.1, …, 0.8 and different *p* are used. The lines show the mean accuracy, and the edges of the envelopes show ±1.96 standard errors (SE). The results are based on 1000simulated datasets where each simulated dataset has 50 replicates.

The general pattern observed in Fig 1 is that log yields the worst results, log-TSS is an improvement, CLR and SparCEV outperform log-TSS (except when sparsity is badly violated), and SparCEV outperforms CLR at low *p*. This behavior is consistent with the theory presented in S1 Text. Situations where *p* is small may be encountered in practice, for example, when abundances at high taxonomic levels are considered or when synthetic communities are employed, as is sometimes done in the plant field [25].

### The Effect of Diversity

Friedman and Alm [13] showed that the accuracy of correlation estimates in case A depends on the diversity of the microbiome. They show that the accuracy of empirical Pearson correlation estimates decreases as *p*_eff_ increases, whereas SparCC is unaffected by *p*_eff_. Fig 2 shows the impact of diversity in case B with *p* = 100 and different average *p*_eff_. The simulation settings are identical to those in Fig 1 for *p* = 100, except that we choose 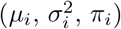 differently (see Parameter Selection under Material and Methods).Specifically, we set *π*_*i*_ = 0 for all *i* to avoid zero inflation. This is because a more zero-inflated dataset will tend to have lower entropy (and thus lower *p*_eff_) than a less zero-inflated dataset. This introduces a chaotic element to the simulation process that may muddle the patterns we seek to investigate.

In Fig 1, we show two sets of lines for the cluster method, one for correlated pairs and one for uncorrelated pairs. In Fig 2, these lines would have fallen on top of each other, so for ease of presentation, the lines for the uncorrelated pairs have been omitted. The results for uncorrelated pairs are instead shown in S2 Fig, where the overall pattern is similar to Fig 2. However, for uncorrelated pairs, both log and log-TSS perform better than CLR and SparCEV when *p*_eff_ is high and sparsity is violated.

In Fig 2, we see that SparCEV and CLR are both only mildly affected by the effective number of OTUs for correlation matrices generated by both the cluster method and the loadings method, regardless of threshold or density of the correlation matrix. SparCEV still consistently outperforms CLR, although the difference is negligible (for all *p*_eff_, the difference is similar to the difference we saw in Fig 1 at *p* = 100). However, the accuracy of the results obtained from log and log-TSS depend heavily on *p*_eff_. The accuracy of log-TSS is similar to that of CLR and SparCEV only for dense correlation matrices with uniformly distributed abundances, which are unlikely to occur in nature. In general, the benefit of using CLR or SparCEV is greater for less diverse microbiota. This is consistent with established knowledge in case A [13].

### Application on Atopic Dermatitis Data

In this section, we analyze the correlations found in an atopic dermatitis dataset from Byrd et al. [31]. The severity of the symptoms was quantified by the widely used measure *objective SCORing of Atopic Dermatitis* (objective SCORAD) [38]. We estimate the correlations between objective SCORAD and bacterial abundances at the family level using SparCEV. We obtained a correlation threshold of 0.52 using the threshold selection approach described in Materials and Methods. Additionally, we calculated empirical bootstrap confidence intervals (CI) with the BC_a_-method according to Efron [39]. The families with absolute correlation with SCORAD exceeding 0.52 are shown in Fig 3A with the exception of two families whose 95%-CIs included zero. These families were *Nocardioidaceae* (Estimate: -0.54, 95%-CI: [-0.76, 0.03]) and *Nakamurellaceae* (Estimate: -0.55, 95%-CI: [-0.77, 0.02]).

It is well known that colonization by *Staphylococcus aureus* can exacerbate the severity of atopic dermatitis [31]. Indeed, we find that *Staphylococcaceae* is the family whose abundance has the strongest positive correlation with the objective SCORAD (estimate: 0.65, 95%-CI: [0.34, 0.81]), see Fig 3A and Fig 3B. Some members of the fungal family *Malasseziaceae* are believed to play a pathogenic role in atopic dermatitis [40, 41], and this family also appears to be positively correlated with the objective SCORAD, although it is not above our permutation threshold (estimate: 0.42, 95%-CI: [0.08, 0.74]). Other studies found that the relative abundance of the genus *Propionibacterium* was depleted in patients with atopic dermatitis [42] and that the genus *Cutibacterium* may inhibit the growth of *Staphylococcus aureus* [43]. Both these genera are members of the family *Propionibacteriaceae*, but we did not find a correlation between the objective SCORAD and the abundance of this family (estimate: -0.20, 95%-CI: [-0.56, 0.20]). The strongest negative correlation detected was with the family *Hyphomicrobiaceae* (estimate: -0.73, 95%-CI: [-0.84, -0.51]), but to our knowledge, this family is not known to play a role in atopic dermatitis. According to Byrd et al., the two predominant viruses were polyomaviruses and papillomaviruses. The Byrd et al. data indicate a possible negative correlation with *Polyomaviridae* (estimate: -0.41, 95%-CI: [-0.71, -0.04]), albeit below our threshold, while *Papillomaviridae* is uncorrelated with SCORAD (estimate: -0.16, 95%-CI: [-0.62, 0.22]). All correlations and their confidence intervals can be found in S1 Table.

Fig 3C shows that the diversity (as measured by effective number of families) is negatively correlated with the objective SCORAD score. This is consistent with prior knowledge that the diversity of the skin microbiome is substantially reduced in atopic dermatitis patients [42, 43]. The effective number of families is only 18 (out of 407 observed families) even in the most diverse replicate (the effective number of families in the least diverse replicate is 1.2, with over 96% of the relative abundance occupied by *Staphylococcaceae*). Thus, the diversity in all replicates is low, and by Fig 2 we expect substantially more accurate correlation estimates from SparCEV or CLR compared with log-TSS. According to Fig 3D, the estimates using log-TSS are consistently smaller than those of SparCEV. Assuming the SparCEV estimates are more accurate, as is indicated by Fig 1 and Fig 2, the log-TSS estimates possibly mask potential positive correlations and exaggerate negative correlations.

### Case C

We repeated the numerical studies from Fig 1 in case C. The left column in Fig 4 shows results with the correlation matrices obtained using the cluster method with *c* = 0.1. For results with other values of *c*, see S4 Fig. On Fig 4, all tested methods perform similarly on non-correlated pairs and on correlated pairs; CLR and CLR+VST yield almost identical results and outperform log-TSS, while SparXCC is superior when *p* or *q* is small, in agreement with the theory. On S4 Fig, we see that for *c* = 0.4, the performance lead for SparXCC is reduced and for *c* = 0.7 it is almost nonexistent. Just as in case B, a non-sparse correlation matrix is a considerable obstacle for SparXCC, CLR, and CLR+VST. They are all outcompeted by log-TSS in this setting. With the loadings method, we see a similar pattern as in case B. When *p* = *q* = 10, SparXCC outcompetes all alternatives, especially for higher thresholds. For *p* = *q* = 100 the difference between SparXCC and CLR or CLR+VST is reduced and for *p* = *q* = 1000, SparXCC, CLR, and CLR+VST all perform identically and all outperform log-TSS. In S5 Fig, we see results for all tested combinations of *p* and *q*. Here, we see that SparXCC outperforms all tested transformation-based methods when either *p* or *q* is sufficiently small. We also carried out the simulations without zero-inflation and found practically identical results to those seen on Fig 4. We note that situations where *p, q* or both are small may arise for example when examining correlations between 16S data (bacterial OTUs) and ITS data (fungal OTUs). Specifically, when synthetic communities are employed or when correlations at a high taxonomic level are of interest.

In case B, SparCEV did not have any meaningful drawbacks in relation to CLR. There are cases where they are practically identical, and then there are cases where SparCEV outcompetes CLR. It is not as simple in case C, since SparXCC requires considerably more computational resources than SparCEV. As a result, it is advisable to carefully consider one’s specific dataset before choosing a method. When both *p* and *q* are large, there is little gain in using SparXCC over CLR, but it runs substantially slower.

### Application on plant microbiome data

In this section, we analyze the correlations found in the root microbiome of *Lotus japonicus* in a dataset by Thiergart et al. [20]. We have two sequencing datasets (each compositional), one from 16S ribosomal RNA (rRNA) and one from internal transcribed spacers (ITS). The 16S data contains bacterial OTUs, and the ITS data contains fungal OTUs. The data are from plants of multiple genotypes, the wild type (Gifu) and the mutants *ccamk, symrk, ram1*, and *nfr5*. The data contains 15–22 replicates for each genotype. We estimate the correlations using SparXCC. The replicates within each genotype originate from three different experiments. This potentially has a confounding effect on the results. For the purposes of this example, we employ the function `RemoveBatcheffect` from the R-package `limma` [44] to correct for differing means between experiments. The results can be seen on Fig 5. More details on correlated OTUs can be found in S2 Table. A similar analysis was carried out on data collected from the rhizosphere of the plant. The results of this can be found in S7 Fig and S3 Table.

**Fig 5.**
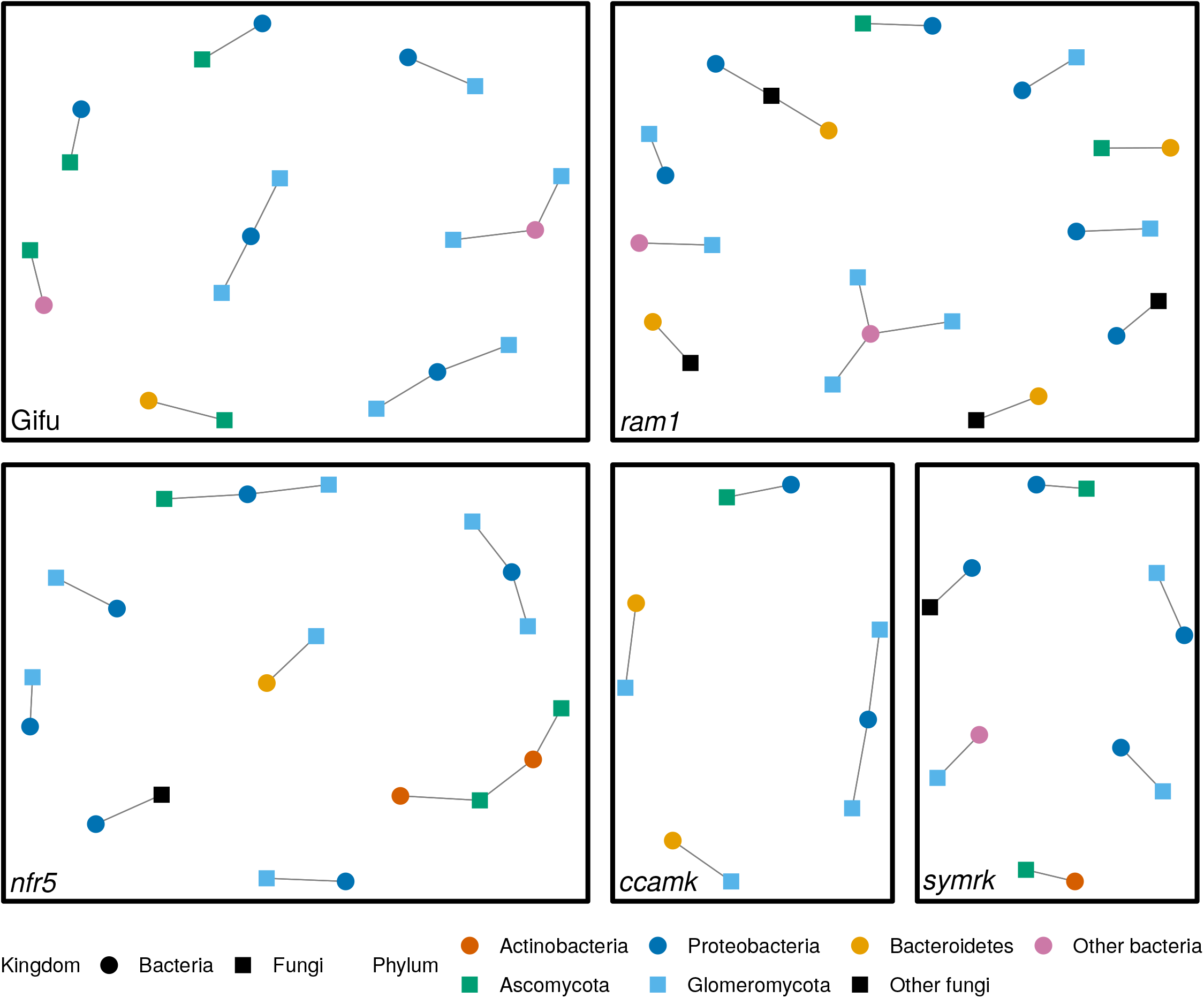
Results from applying CLR to 16S and ITS sequencing data from the root microbiome of *Lotus japonicus*, from Thiergart et al. [20]. Each circular vertex represents a bacterial OTU from the 16S data and a square vertex represents a fungal OTU from the ITS data. Vertices are colored based on the phylum of the OTU it represents. Two vertices are connected by an edge if their estimated correlation is above the permutation threshold. The analysis is carried out separately for the genotypes Gifu, *ram1, nfr5, ccamk*, and *symrk*. Only cross-correlations are shown.

Thiergart et al. estimate cross-correlations between bacterial and fungal OTUs only on the Gifu data using Spearman correlations on TSS-transformed data (Spearman-TSS). They consider a pair correlated when the *p*-value is less than 0.001 and thus obtain 585 pairs with significant correlations. Using permutation thresholding on SparXCC, we find only 12 correlated pairs in Gifu when correcting for confounding effects. In order to make a direct comparison to the results in the original paper, we also carried out the correlation estimation without correcting for confounding effects. We then obtain 1229 correlations above the threshold. A substantial proportion of correlations identified by Thiergart et. al were also found by SparXCC (66 %). On S6 Fig and in S4 Table, we see that in most cases, the methods find similar estimates, but in some cases they may differ considerably. In fact, in some cases, the two methods disagree on the sign of the correlation. The reason for the differences between the methods may be that SparXCC approximates Pearson correlations, which measure linear relationships, while Spearman correlations measure monotonic relationships. To examine this possibility, we also estimated the Pearson correlations of the log-TSS transformed data (Pearson-log-TSS) and found 1395 pairs with significant correlations. Interestingly, we found that Pearson-log-TSS showed a greater degree of overlap with Spearman-TSS than with SparXCC (87 % vs 72 %).

Of the pairs where SparXCC and Spearman-TSS disagreed on the sign, 21 were above the permutation threshold, but not detected as significant by the *t* -test. All of these pairs involved two specific fungal OTUs, both members of the phylum Ascomycota. Both had many reads (ranging from 95 to 2576), so these results are not an artifact of low read counts. Additionally, all of these pairs showed the same pattern when comparing SparXCC to Pearson-log-TSS; the estimated correlations had different signs, but were not detected as significant by the *t* -test. This suggests that the model structure in SparXCC is able to capture some pattern of association that is lost with the transformation-based methods.

## Discussion

For the theoretical considerations in this paper, we, like Friedman and Alm [13], assume that the data follow the model in (1). According to this model, the true relative abundances *r*_*i*_ are observed, which would only be the case with infinite sequencing depth. We nevertheless assess the different correlation estimation methods using data simulated under a more realistic setting where the *x*_*i*_s are noisy observations of the *r*_*i*_s. Specifically, SparseDOSSA2 assumes that the *x*_*i*_s are multinomial, given the library size *N* and the *r*_*i*_s. Friedman and Alm [13] suggests mitigating the impact of the technical variance of *x*_*i*_ given (*r*_*i*_, *N*) by using a Monte Carlo sampling procedure based on a uniform Dirichlet prior. However, in our simulation setup, we find that this reduces accuracy compared to using a pseudo-count (See Fig S8 Fig). It is a topic of further research to investigate the nature of the technical variance (e.g. if it is truly generated by a multinomial model, as postulated by SparDossa2) and how to account for it in the cross-correlation estimations.

We did not consider testing null hypotheses of zero correlation. Due to various sources of bias, including the aforementioned technical variance, it is difficult to base hypothesis testing on theoretical results. Friedman and Alm [13] use a bootstrapping procedure when applying SparCC in case A. This is computationally demanding, however, not least since corrections for multiple testing are needed when carrying out hypothesis testing for a large number of correlations. Furthermore, in cases B and C, it is challenging to construct a bootstrap simulation scheme that respects the null hypothesis for a particular correlation while maintaining the remaining correlation structure. Due to these difficulties, we believe that it may be more appropriate to rely on the correlation estimates themselves, as we have done with the permutation threshold selection.

A fundamental assumption in this paper is that the interactions between microbes and other variables can be adequately described by a correlation matrix. To our knowledge, no alternatives have been unambiguously shown to universally better describe interactions between compositional datasets such as microbiome and RNA-seq data. Which metric is more sensible may depend on the underlying biology of the specific data under study. We compared the performance of SPIEC-EASI and correlation-based approaches in case C. SPIEC-EASI uses a penalized regression scheme to estimate the precision matrix, *R*^-1^, and does not aim to estimate the correlation matrix, *R*, directly. Instead the primary aim is to discriminate between pairs that are conditionally independent and pairs that are not. With a correlation matrix constructed using the cluster method, a pair is uncorrelated if and only if it is conditionally independent 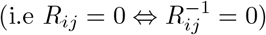. Thus we can directly compare the power and false discovery rate (FDR) of SPIEC-EASI with those found using pair-wise correlations.To do this, we subjected the correlation estimates of CLR to a *t* -test and the estimates of SparXCC to the permutation thresholding scheme described in the material and methods section as well as using SPIEC-EASI to identify conditionally dependent pairs. Compared to the thresholding method, we found that SPIEC-EASI has higher power for *n ≤* 50 but with a much higher FDR. The *t* -test had better power than the other methods tested for *n ≥* 50, but FDR increased as *n* increased. However, it was able to adequately control the FDR at *n ≤*50. Permutation thresholding saw relatively high FDR at *n* = 20, but was otherwise able to better control the FDR than the other methods tested, and it was nearly as powerful as the *t*-test. See S10 Fig for similar simulations in case B, comparing a *t* -test, permutation thresholding with *m* as a threshold, and permutation thresholding with *m*\* as a threshold.

The interactions present in a real biological system are likely to be more complicated than a correlation matrix generated by the cluster method can account for. In such cases, the methods may diverge (in general, 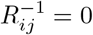 need not imply that *R*_*ij*_ = 0 or vice versa), and it may not be clear which is more appropriate.

## Conclusion

When estimating correlations between compositional variables and non-compositional variables (case B), the results in Fig 1, Fig 2, and S1 Fig suggest that SparCEV should be the method of choice. When the number of compositional variables is sufficiently large (above 100 in our results), empirical Pearson correlations following the CLR-transformation perform essentially just as well. When estimating cross-correlations between two compositional datasets (case C), the results in Fig 4, S3 Fig, S4 Fig, and S5 Fig suggest that the method of choice should be SparXCC. However, when both *p* and *q* are large, CLR may be preferred, since it performs essentially just as well in these cases, but with considerably less computational complexity.

## Supporting information

**S1 Fig. Case B without biological zeros** Accuracy of the different cross-correlation methods in case B, in the absence of biological zero by enforcing *π*_*j*_ = 0 for *j* = 1, …, *p*. Otherwise, the same simulation settings as Fig 1 are used.

**S2 Fig. Case B diversity and zero correlations** Accuracy of the different cross-correlation methods in case B on uncorrelated pairs at different levels of diversity.

**S3 Fig. Case C without biological zeros** Accuracy of the different cross-correlation methods in case C, in the absence of biological zero by enforcing *π*_*j*_ = 0 for *j* = 1, …, *p* + *q*. Otherwise, the same simulation settings as Fig 4 are used.

**S4 Fig. Cluster method in case C for large *c*** Accuracy of the different cross-correlation methods on correlation matrices generated by the cluster method in case C for *c* = 04, 0.7. Otherwise, the same simulation settings as Fig 4 are used.

**S5 Fig. Loadings method in case C for all combinations of *p* and *q*** Accuracy of the different cross-correlation methods on correlation matrices generated by the loadings method in case C for all combinations of *p* = 10, 100, 1000 and *q* = 10, 100, 1000. Otherwise, the same simulation settings as Fig 4 are used.

**S6 Fig. Spearman correlations of relative abundances vs SparXCC** The estimated correlation coefficients as estimated by Spearman correlations of relative abundances plotted against correlations approximated by SparXCC. For Spearman, a pair is considered correlated when a *t* -test returns a *p*-value less than 0.001. For SparXCC, a pair is considered correlated when it is above the permutation threshold.

**S7 Fig. Cross-correlation network constructed on rhizosphere data** Graph with edges between nodes when the cross-correlation is above a permutation threshold, estimated by SparXCC on rhizosphere data.

**S8 Fig. Pseudo-count versus Dirichlet Monte Carlo sampling** Accuracy of using a pseudo-count versus Dirichlet Monte Carlo for SparCEV.

**S9 Fig. Separating correlated and uncorrelated pairs in Case C** Power and FDR of CLR with a *t* -test, SparXCC with permutation thresholding, and SPIEC-EASI.

**S10 Fig. Separating correlated and uncorrelated pairs in Case B** Power and FDR of CLR with a *t* -test and SparCEV with permutation thresholding, using both *m* and *m*\* as defined in section Discriminating between Correlated and Uncorrelated pairs.

**S1 Table** Correlations between families and objective SCORAD score.

**S2 Table** Correlations between bacterial OTUs from 16S data and fungal OTUs from ITS data from the root of *Lotus japonicus*. Confounding effects from the experiment effect were removed and SparXCC was applied.

**S3 Table** Correlations between bacterial OTUs from 16S data and fungal OTUs from ITS data from the rhizosphere of *Lotus japonicus*. Confounding effects from the experiment effect were removed and SparXCC was applied.

**S4 Table** Correlations between bacterial OTUs from 16S data and fungal OTUs from ITS data from the root of *Lotus japonicus*. The data was not corrected for confounding effects prior to correlation estimation.

**S1 Text** Theoretical analysis of transformation-based correlations, derivation of compositionally aware methods, and construction of correlation matrices

