## Supplementary material for "Compositionally aware estimation of cross-correlations for microbiome data": S1 text

Throughout this supplementary note, we use the same notation as in the main text of the paper.

### 1 Transformation-based Correlation Approximations

We have

$$\mathbb{C}\text{ov}\left[\log x_i, b\right] = \mathbb{C}\text{ov}\left[\log a_i, b\right] - \mathbb{C}\text{ov}\left[\log A, b\right] + \mathbb{C}\text{ov}\left[\log N, b\right] \approx \mathbb{C}\text{ov}\left[\log a_i, b\right],$$

provided the phenotype variable is approximately uncorrelated with the library size and the total bacterial load. With similar assumptions, we have

$$\mathbb{C}\text{ov}\left[\log \text{TSS}(x_i), b\right] = \mathbb{C}\text{ov}\left[\log a_i, b\right] - \mathbb{C}\text{ov}\left[\log A, b\right] \approx \mathbb{C}\text{ov}\left[\log a_i, b\right].$$

Finally, we also have

$$\begin{aligned}\mathbb{C}\text{ov}\left[\text{CLR}(x_i), b\right] = \mathbb{C}\text{ov}\left[\text{CLR}(a_i), b\right] &= \mathbb{C}\text{ov}\left[\log a_i, b\right] - \frac{1}{p}\sum_{j=1}^{p}\mathbb{C}\text{ov}\left[\log a_j, b\right] \\ &\approx \mathbb{C}\text{ov}\left[\log a_i, b\right],\end{aligned}$$

which holds, when $\frac{1}{p}\sum_{j=1}^{p}\mathbb{C}\text{ov}\left[\log a_j, b\right] \approx 0$. Thus, we see that all approaches can approximate $\mathbb{C}\text{ov}\left[\log a_i, b\right]$ under appropriate assumptions. For the variance, we have

$$\begin{aligned}\mathbb{V}\text{ar}\left[\log x_i\right] &= \mathbb{V}\text{ar}\left[\log a_i\right] + \mathbb{V}\text{ar}\left[\log A\right] - 2\mathbb{C}\text{ov}\left[\log a_i, \log A\right] + \mathbb{V}\text{ar}\left[\log N\right] \\ &\approx \mathbb{V}\text{ar}\left[\log a_i\right] + \mathbb{V}\text{ar}\left[\log A\right] + \mathbb{V}\text{ar}\left[\log N\right].\end{aligned}$$

Note that the above approximation holds when $\mathbb{C}\text{ov}\left[\log a_i, \log A\right] \approx 0$. This may be violated, if OTU $i$ makes up a sufficiently large proportion of the microbiome or if it is strongly correlated with many other OTUs. Regardless, we cannot assume that $\mathbb{V}\text{ar}\left[\log A\right]$ or $\mathbb{V}\text{ar}\left[\log N\right]$ are small, and thus we do not get a good approximation of $\mathbb{V}\text{ar}\left[\log a_i\right]$ using log-transformed read-counts. We have

$$\begin{aligned}\mathbb{V}\text{ar}\left[\log \text{TSS}(x_i)\right] &= \mathbb{V}\text{ar}\left[\log a_i\right] + \mathbb{V}\text{ar}\left[\log A\right] - 2\mathbb{C}\text{ov}\left[\log a_i, \log A\right] \\ &\approx \mathbb{V}\text{ar}\left[\log a_i\right] + \mathbb{V}\text{ar}\left[\log A\right],\end{aligned}$$

and thus we do not get a good approximation of $\mathbb{V}\text{ar}\left[\log a_i\right]$ from log-TSS either. Finally, we have,

$$\begin{aligned}\mathbb{V}\text{ar}\left[\text{CLR}(x_i)\right] &= \mathbb{V}\text{ar}\left[\log a_i\right] + \mathbb{V}\text{ar}\left[\frac{1}{p}\sum_{j=1}^{p}\log a_j\right] - 2\mathbb{C}\text{ov}\left[\log a_i, \frac{1}{p}\sum_{j=1}^{p}\log a_j\right] \\ &= \mathbb{V}\text{ar}\left[\log a_i\right] + \frac{1}{p^2}\sum_{j=1}^{p}\sum_{k=1}^{p}\mathbb{C}\text{ov}\left[\log a_j, \log a_k\right] - \frac{1}{p}\sum_{j=1}^{p}2\mathbb{C}\text{ov}\left[\log a_i, \log a_j\right] \\ &\approx \frac{p-2}{p}\mathbb{V}\text{ar}\left[\log a_i\right] + \frac{1}{p^2}\sum_{j=1}^{p}\mathbb{V}\text{ar}\left[\log a_j\right] \approx \mathbb{V}\text{ar}\left[\log a_i\right],\end{aligned}$$

where the last relation holds when $p$ is large, and the second to last relation hold when

$$\frac{1}{p^2}\sum_{k=1}^{p}\sum_{j\neq k}\mathbb{C}\mathrm{ov}\left[\log a_j, \log a_k\right] \approx 0, \quad \frac{1}{p}\sum_{j\neq i}\mathbb{C}\mathrm{ov}\left[\log a_i, \log a_j\right] \approx 0.$$

#### 2 Derivation of SparCEV

Suppose $x_i = a_i / \sum_j a_j$ for absolute abundances $a_1,\ldots,a_p$ and that $y$ is some scalar nominal variable. Then,

$$\mathbb{C}\mathrm{ov}\left[\log\frac{x_i}{x_j}, y\right] = \mathbb{C}\mathrm{ov}\left[\log\frac{a_i}{a_j}, y\right] = \mathbb{C}\mathrm{ov}\left[\log a_i, y\right] - \mathbb{C}\mathrm{ov}\left[\log a_j, y\right].$$

Thus

$$\sum_{j\neq i}\mathbb{C}\mathrm{ov}\left[\log\frac{x_i}{x_j}, y\right] = (p-1)\left(\mathbb{C}\mathrm{ov}\left[\log a_i, y\right] - \frac{1}{p-1}\sum_{j\neq i}\mathbb{C}\mathrm{ov}\left[\log a_j, y\right]\right) \approx (p-1)\mathbb{C}\mathrm{ov}\left[\log a_i, y\right],$$

where the approximation comes from the assumption that $\frac{1}{p-1}\sum_{j\neq i}\mathbb{C}\mathrm{ov}\left[\log a_j, y\right] \approx 0$. Thus we get

$$\mathbb{C}\mathrm{orr}\left[\log a_i, y\right] \approx \frac{1}{\sigma_y \alpha_i}\frac{1}{p-1}\sum_{j\neq i}\mathbb{C}\mathrm{ov}\left[\log\frac{x_i}{x_j}, y\right],$$

where $\sigma_y^2 = \mathbb{V}\mathrm{ar}\left[y\right]$, and $\alpha_i^2 = \mathbb{V}\mathrm{ar}\left[\log a_i\right]$. The variance $\sigma_y^2$ can be estimated in a standard fashion and $\alpha_i^2$ can be estimated using SparCC assuming that most abundances are not correlated.

#### 3 Derivation of SparXCC

Suppose we have absolute abundances $a = (a_1, \ldots, a_p)^\top$, and absolute gene expression levels $b = (b_1, \ldots, b_q)^\top$. Let $N_a = a^\top 1_p$ and $N_b = b^\top 1_q$ and define the relative features $x = a/N_a$ and $y = b/N_b$. We have $a_i = N_a x_i$ and $b_k = N_b y_k$ and we seek the cross-correlations between variables $a_i$ and $b_k$. First note that

$$\begin{aligned}
\sum_{j=1}^{p}\mathbb{V}\mathrm{ar}\left[\log\frac{x_i}{x_j}\right] = \sum_{j=1}^{p}\mathbb{V}\mathrm{ar}\left[\log\frac{a_i}{a_j}\right] &= p\alpha_i^2 + \sum_{j=1}^{p}\alpha_j^2 - 2\sum_{j=1}^{p}\alpha_i\alpha_j\rho_{ij} \\
&= (p-1)\alpha_i^2 + \sum_{j\neq i}\alpha_j^2 - 2\sum_{j\neq i}\alpha_i\alpha_j\rho_{ij} \\
&= (p-1)\alpha_i^2\left(1 + \frac{1}{p-1}\sum_{j\neq i}\frac{\alpha_j^2}{\alpha_i^2} - 2\frac{1}{p-1}\sum_{j\neq i}\frac{\alpha_j}{\alpha_i}\rho_{ij}\right) \\
&\approx (p-1)\alpha_i^2 + \sum_{j\neq i}\alpha_j^2, \qquad (1)
\end{aligned}$$

where $\alpha_i^2 = \mathbb{V}\mathrm{ar}\left[\log a_i\right]$ and the last equality follows from the assumption that

$$1 + \frac{1}{p-1}\sum_{j\neq i}\frac{\alpha_j^2}{\alpha_i^2} \gg 2\frac{1}{p-1}\sum_{j\neq i}\frac{\alpha_j}{\alpha_i}\rho_{ij}.$$

In the case where $\alpha_i^2 = \alpha_j^2$ for all $i$, $j$, this reduces to the assumption that

$$1 \gg \frac{1}{p-1}\sum_{j\neq i}\rho_{ij},$$

that is, the average correlation between OTU $i$ and another OTU is small. Under similar assumptions, we have

$$\mathbb{V}\text{ar}\left[\log\frac{y_k}{y_l}\right] \approx (q-1)\beta_k^2 + \sum_{l\neq k}\beta_l^2, \tag{2}$$

where $\mathbb{V}\text{ar}\left[\log b_k\right] = \beta_k^2$. SparCC applies (1) and (2) to approximate the variances $\alpha_i^2$ and $\beta_k^2$ for $i = 1, \ldots, p$ and $j = 1, \ldots, q$.

When analysing compositional data, it is common to consider ratios as above, but

$$\log\frac{a_i}{b_k} = \log\frac{N_a x_i}{N_b y_k} = \log\frac{x_i}{y_k} + \log\frac{N_a}{N_b}$$

where $N_a$ and $N_b$ are unobserved. Instead, we may consider ratios of ratios. Specifically,

$$\log\frac{a_i}{b_k} - \log\frac{a_j}{b_l} = \left(\log\frac{x_i}{y_k} + \log\frac{N_a}{N_b}\right) - \left(\log\frac{x_j}{y_l} + \log\frac{N_a}{N_b}\right) = \log\frac{x_i}{y_k} - \log\frac{x_j}{y_l}.$$

Thus,

$$t_{ijkl} := \mathbb{V}\text{ar}\left[\log\frac{a_i}{b_k} - \log\frac{a_j}{b_l}\right] = \mathbb{V}\text{ar}\left[\log\frac{a_i}{b_k}\right] + \mathbb{V}\text{ar}\left[\log\frac{a_j}{b_l}\right] - 2\mathbb{C}\text{ov}\left[\log\frac{a_i}{b_k}, \log\frac{a_j}{b_l}\right]. \tag{3}$$

We now take the sum $t_{ik} := \sum_{j=1}^{p}\sum_{l=1}^{q} t_{ijkl}$ and by (1) and (2) we obtain

$$t_{ik} \approx (p-1)q\alpha_i^2 + q\sum_{j\neq i}\alpha_j^2 + p(q-1)\beta_k^2 + p\sum_{l\neq k}\beta_l^2 - 2\sum_{j=1}^{p}\sum_{l=1}^{q}\mathbb{C}\text{ov}\left[\log\frac{a_i}{b_k}, \log\frac{a_j}{b_l}\right]. \tag{4}$$

For the last term in (4), we obtain

$$\begin{aligned}
\sum_{j=1}^{p}\sum_{l=1}^{q}\mathbb{C}\text{ov}\left[\log\frac{a_i}{b_k}, \log\frac{a_j}{b_l}\right] =& pq\mathbb{C}\text{ov}\left[\log a_i, \log b_k\right] - q\sum_{j=1}^{p}\mathbb{C}\text{ov}\left[\log a_j, \log b_k\right] \\
&- p\sum_{l=1}^{q}\mathbb{C}\text{ov}\left[\log a_i, \log b_l\right] + \sum_{j=1}^{p}\sum_{l=1}^{q}\mathbb{C}\text{ov}\left[\log a_j, \log b_l\right] \\
=& (p-1)(q-1)\mathbb{C}\text{ov}\left[\log a_i, \log b_k\right] - (q-1)\sum_{j\neq i}\mathbb{C}\text{ov}\left[\log a_j, \log b_k\right] \\
&- (p-1)\sum_{l\neq k}\mathbb{C}\text{ov}\left[\log a_i, \log b_l\right] + \sum_{j\neq i}\sum_{l\neq k}\mathbb{C}\text{ov}\left[\log a_j, \log b_l\right] \\
=& (p-1)(q-1)\alpha_i\beta_k\rho_{ik} - (q-1)\sum_{j\neq i}\alpha_j\beta_k\rho_{jk} \\
&- (p-1)\sum_{l\neq k}\alpha_i\beta_l\rho_{il} + \sum_{j\neq i}\sum_{l\neq k}\alpha_j\beta_l\rho_{jl} \\
=& (p-1)(q-1)\alpha_i\beta_k\Bigg(\rho_{ik} - \frac{1}{p-1}\sum_{j\neq i}\frac{\alpha_j}{\alpha_i}\rho_{jk} \\
&- \frac{1}{q-1}\sum_{l\neq k}\frac{\beta_l}{\beta_k}\rho_{il} + \frac{1}{(p-1)(q-1)}\sum_{j\neq i}\sum_{l\neq k}\frac{\alpha_j}{\alpha_i}\frac{\beta_l}{\beta_k}\rho_{jl}\Bigg) \\
\approx& (p-1)(q-1)\alpha_i\beta_k\rho_{ik},
\end{aligned}$$

where the approximation follows from the assumption that the last three terms are small compared to the first four terms in (4). Inserting the result above into (4) yields

$$t_{ik} \approx (p-1)q\alpha_i^2 + q\sum_{j\neq i}\alpha_j^2 + p(q-1)\beta_k^2 + p\sum_{l\neq k}\beta_l^2 - 2(p-1)(q-1)\alpha_i\beta_k\rho_{ik}.$$

By re-arranging this, we obtain

$$\rho_{ik} \approx \frac{(p-1)q\alpha_i^2 + p(q-1)\beta_k^2 + q\sum_{j\neq i}\alpha_j^2 + p\sum_{l\neq k}\beta_l^2 - t_{ik}}{2(p-1)(q-1)\alpha_i\beta_k}. \quad (5)$$

Thus, we estimate

$$\hat{\rho}_{ik} = \frac{(p-1)q\hat{\alpha}_i^2 + p(q-1)\hat{\beta}_k^2 + q\sum_{j\neq i}\hat{\alpha}_j^2 + p\sum_{l\neq k}\hat{\beta}_l^2 - \hat{t}_{ik}}{2(p-1)(q-1)\hat{\alpha}_i\hat{\beta}_k}. \quad (6)$$

##### Efficient Estimation of Variance Sums

Exploiting symmetry $t_{ijkl} = t_{jilk}$ and that $t_{iikk} = 0$, a naive approach to estimating the $t_{ik}$s involves estimation of $pq(pq-1)/2$ variances $t_{ijkl} = \mathbb{V}\mathrm{ar}\left[\log\frac{x_i}{y_k} - \log\frac{x_j}{y_l}\right]$. In practice this may be not be feasible since $p$ and $q$ are often fairly large, e.g $p = 5779$ and $q = 3360$ in the genotype data analyzed in the paper. We would then need to estimate approximately 188 trillion variances. Using 32-bit number representations this would take up 685 TB of memory and be prohibitively time consuming. Fortunately it is possible to estimate the $t_{ik}$s without estimating each term in the sum. Recall the expansion

$$\begin{aligned}\mathbb{V}\mathrm{ar}\left[\log\frac{x_i}{y_k} - \log\frac{x_j}{y_l}\right] &= \mathbb{V}\mathrm{ar}\left[\log\frac{x_i}{y_k}\right] + \mathbb{V}\mathrm{ar}\left[\log\frac{x_j}{y_l}\right] - 2\mathbb{C}\mathrm{ov}\left[\log\frac{x_i}{y_k}, \log\frac{x_j}{y_l}\right] \\ &= \mathbb{V}\mathrm{ar}\left[\log\frac{x_i}{y_k}\right] + \mathbb{V}\mathrm{ar}\left[\log x_j\right] + \mathbb{V}\mathrm{ar}\left[\log y_l\right] - 2\mathbb{C}\mathrm{ov}\left[\log x_j, \log y_l\right] - 2\mathbb{C}\mathrm{ov}\left[\log\frac{x_i}{y_k}, \log\frac{x_j}{y_l}\right].\end{aligned}$$

The sum over $j$ and $l$ can be written as

$$\begin{aligned}t_{ik} =& pq\mathbb{V}\mathrm{ar}\left[\log\frac{x_i}{y_k}\right] + q\sum_{j=1}^{p}\mathbb{V}\mathrm{ar}\left[\log x_j\right] + p\sum_{l=1}^{q}\mathbb{V}\mathrm{ar}\left[\log y_l\right] \\ &- 2\mathbb{C}\mathrm{ov}\left[\sum_{j=1}^{p}\log x_j, \sum_{l=1}^{q}\log y_l\right] - 2\mathbb{C}\mathrm{ov}\left[\log\frac{x_i}{y_k}, q\sum_{j=1}^{p}\log x_j - p\sum_{l=1}^{q}\log y_l\right]. \end{aligned} \quad (7)$$

Thus estimating all $t_{ik}$s by estimating each term in (7) requires only $2pq + p + q + 1$ estimates. In the example above, this would require only about 150 MB of memory, and can be carried out in a reasonable time frame.

#### 4 Construction of correlation matrices

We consider two approaches to construct the correlation matrix $R$ needed for the simulation scheme detailed above.

##### Cluster Method

Suppose we have OTU abundances $a_1, \ldots, a_p$ and other variables $b_{p+1}, \ldots, b_{p+q}$. A correlation matrix $R$ is constructed as follows.

1. Pick a proportion $0 < c < 1$.
2. Randomly select a set of indices, $M_a \subseteq \{1, 2, \ldots, p\}$ containing $\max\{\lfloor cp\rfloor, 1\}$ indices and a set $M_b \subseteq \{1, 2, \ldots, q\}$ containing $\max\{\lfloor cq\rfloor, 1\}$ indices.
3. Simulate $c^+ \sim \mathrm{Unif}(0, 1)$
4. Let $M_a^+$ be a set of $\lfloor c^+|M_a|\rfloor$ indices, sampled without replacement from $M_a$ and let $M_a^- = M_a \backslash M_a^+$. Construct $M_b^+$ and $M_b^-$ in the same way.

5. Let $M = M_a \cup M_b$, $M^+ = M_a^+ \cup M_b^+$ and $M^- = M_a^- \cup M_b^-$.

6. Pick a correlation strength, $\rho \in (0, 1)$.

7. Construct $R$ with ones on the diagonal and off-diagonal entries according to Table 1.

It is easy to check that the generated $R$ is positive definite. The interpretation of the procedure above is that we construct a cluster of OTUs and a cluster of other variables. Non-zero positive or negative correlation exists within and between clusters. Variables outside the clusters are not correlated with any other variables. This produces a simple correlation matrix where it is easy to control the level of sparsity and the strength of the correlations.

| | $j \in M^+$ | $j \in M^-$ | $j \notin M$ |
|---|---|---|---|
| $i \in M^+$ | $\rho$ | $-\rho$ | 0 |
| $i \in M^-$ | $-\rho$ | $\rho$ | 0 |
| $i \notin M$ | 0 | 0 | 0 |

Table 1: Off-diagonal entries $R_{ij}$ of the matrix constructed with the cluster method.

#### Loadings Method

In addition to the cluster method, we also employ a procedure that does not produce zero correlations but where correlations are generally small with a few outlying large correlations.

1. Choose some $k < p + q$ and generate a $p + q \times k$ matrix $Q$ with independent random entries (here generated from a standard normal distribution).

2. Let $U = QQ^\top$. $U$ is symmetric and positive semi-definite, but rank-deficient.

3. Generate a $p + q \times p + q$ diagonal matrix with a positive diagonal (here generated as absolute values of independent standard normal variables).

4. Let $C = U + D$. $C$ is a symmetric full-rank positive definite matrix, and thus the covariance matrix of a non-degenerate distribution.

5. Let $R = E^{-1/2} C E^{-1/2}$, where $E = \text{diag}\{C_{11}, \ldots, C_{p+q,p+q}\}$.

The above procedure does not guarantee that the sparsity assumption holds. However, in practice we find that the average correlations are typically close to 0, provided $p + q$ is not too small. For large $k$ the correlations are more concentrated around 0. Since we want presence of highly correlated pairs of variables we consider $k = 5$.
